# Engineering Lipid Droplet Assembly Mechanisms for Improved Triacylglycerols Accumulation in *S. cerevisiae*

**DOI:** 10.1101/261057

**Authors:** Paulo Gonçalves Teixeira, Florian David, Verena Siewers, Jens Nielsen

## Abstract

Production of triacylglycerols (TAGs) through microbial fermentation is an emerging alternative to plant and animal-derived sources. The yeast *Saccharomyces cerevisiae* is a preferred organism for industrial use but has natively a very poor capacity of TAG production and storage. Here, we engineered *S. cerevisiae* for accumulation of high TAG levels through the use of structural and physiological factors that influence assembly and biogenesis of lipid droplets. First, human and fungal perilipin genes were expressed, increasing TAG content by up to 36% when expressing the human perilipin gene *PLIN3.* Secondly, expression of the *FIT2* homologue *YFT2* resulted in a 26% increase in TAG content. Lastly, the genes *ERD1* and *PMR1* were deleted in order to induce an ER stress response and stimulate lipid droplet formation, increasing TAG content by 72% for Δ*erd1*, with an additive effect for both *YFT2* and *PLIN3* expression. These new approaches were implemented in previously engineered strains that carry high flux of fatty acid biosynthesis and conversion of acyl-CoA into TAG, resulting in improvements of up to 138% over those high-producing strains without any substantial growth effects or abnormal cell morphology. We find that these approaches are not only a major advancement in engineering *S. cerevisiae* for TAG production, but also highlight the importance of lipid droplet dynamics for high lipid accumulation in yeast.

## 1 Introduction

Production of lipids through microbial fermentation is a progressing alternative to petroleum- and plant-derived sources. Among its advantages are the potential for production of tailored chemical precursors, together with versatility of applications, processes and sustainable sources of raw materials (Pfleger, Gossing and Nielsen 2015). The technology has seen vast progress in the past few years (Zhao *et al.* 2008; Tsigie *et al.* 2011; Galafassi *et al.* 2012) and production of lipids through yeast fermentation is starting to see its light as a viable industrial process.

Yeasts store fatty acids in form of storage lipids, i.e. neutral lipids such as triacylglycerols (TAGs) and sterol esters (SEs) (Czabany, Athenstaedt and Daum 2007) that are accumulated intracellularly in a particular organelle known as the lipid droplet (LD). Lipid droplets have received a lot of attention the past few years and have been consistently progressing from being considered a simple fat reserve towards being recognized as an important intervenient in a variety of cellular functions and having a close relationship with many organelles such as the endoplasmic reticulum (ER), mitochondria, peroxisomes and vacuoles (Beller *et al.* 2010; Brasaemle and Wolins 2012; Schuldiner and Bohnert 2017).

LDs are dynamic organelles with their own life cycle that are formed at the ER and can later mature within the cytoplasm (Wilfling *et al.* 2014; Joshi, Zhang and Prinz 2017). Contained storage lipids can be mobilized by the cell for membrane lipid remodeling or for fatty acid oxidation as an energy and carbon source. Though being in recent scientific focus, the detailed mechanisms involved in the formation and budding of the LD from the ER, its cellular dynamics and the roles of associated proteins are still not completely understood (Welte 2015).

Even though some microbial species are known to accumulate high levels of lipids, the yeast *Saccharomyces cerevisiae* has been the target of many engineering efforts for production of lipid-related species(Zhou *et al.* 2016; d’Espaux *et al.* 2017; Fernandez-Moya and Da Silva 2017), since it is regarded as a robust and well adapted organism for industrial production of a variety of products ranging from pharmaceuticals to chemical building blocks (Krivoruchko and Nielsen 2015; Nielsen and Keasling 2016). *S. cerevisiae* is also a model organism for more fundamental studies of many conserved pathways in eukaryotes. As such, lipid metabolism has been thoroughly studied in this organism and it is known that lipid droplet dynamics are conserved with many other eukaryotes (Petranovic *et al.* 2010; Natter and Kohlwein 2012; Kohlwein, Veenhuis and van der Klei 2013; Welte 2015).

While *S. cerevisiae* natively only accumulates up to 1% of its dry biomass weight as TAGs, it has been successfully engineered to accumulate as much as 25% with 1.7 g/L of TAG produced from 20 g/L of glucose (Ferreira et al., 2018). These engineering efforts have been related to increasing enzyme activity within the TAG synthesis pathway involving mainly upregulation of 2 one flux-limiting enzyme in fatty acid synthesis (the acetyl-CoA carboxylase Acc1), overexpression of enzymes involved in the TAG synthesis pathway (phosphatidate phosphatase Pah1 and diacylglycerol acyltransferase Dga1) and knockout of TAG-hydrolysis pathways (lipases Tgl3/4/5).

Although Pah1 and Dga1 are two important enzymes in LD biogenesis (Barbosa *et al.* 2015; Thiam and Beller 2017), there are a number of other proteins involved in budding of the ER membrane into a lipid droplet structure and later detachment of the droplet structure from the membrane (Brasaemle and Wolins 2012). Pah1, an homologue of the human lipin gene (Han, Wu and Carman 2006), is recruited from the cytosol to the ER membrane through dephosphorylation by the Nem1-Spo7 complex and is responsible for conversion of the phosphatidic acid (PA) existent in the membrane to diacylglycerols (DAGs) (Santos‐Rosa *et al.* 2005; Karanasios *et al.* 2010; Adeyo *et al.* 2011; Barbosa *et al.* 2015). TAGs are then produced from these DAGs by Dga1 or by the acyltransferase Lro1 and are the main component of the lipid droplet core structure. Simultaneously, there are three other main families of proteins known to be directly involved in structural aspects of the LD biogenesis process: seipins, fat storage-inducing transmembrane (FIT) proteins and perilipins (Chen and Goodman 2017). These proteins aid the creation and enable stabilization of a hydrophobic pocket structure containing DAGs and TAGs between the two phospholipid layers of the outside ER membrane called the “nucleation site”, where nascent lipid droplets begin their assembly (Wilfling *et al.* 2013; Thiam and Beller 2017). The nucleation site expands with further accumulation of neutral lipids, consecutively forming a lipid droplet through a protrusion of the external ER membrane (Choudhary *et al.* 2015) (Figure 1).

**Figure 1.**
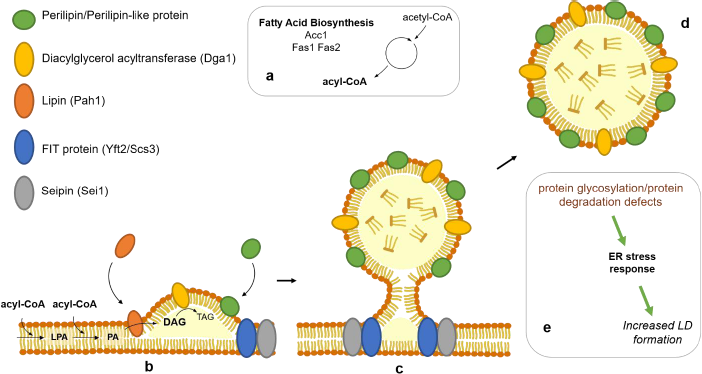
Schematic representation of the different elements involved in lipid droplet formation and budding explored in this study. A) Representation of fatty acid biosynthesis and the 3 genes involved: *ACC1, FAS1* and *FAS2*. Strains used in this study have been engineered for overexpression of the TAG production pathway. An *ACC1* double mutant was expressed to release repression of fatty acid biosynthesis and increase supply of acyl-CoA. PAH1 and DGA1 were overexpressed in order to increase the flux from acyl-CoA to TAGs. b) Nascent lipid droplet assembly encompasses recruitment of Pah1 to the ER membrane and is additional characterized by presence of transmembrane seipins and FIT proteins. Diacylglycerol acyltransferases and some perilipins are also recruited to nascent lipid droplets. c) Seipin is present in LD-ER contact sites and allows for detachment of the LD from the ER. LDs are also matured through incorporation of more fatty acids as TAGs by containing most enzymes in the TAG synthesis pathway. d) Perilipins are usually found coating mature LDs and can be important for maturation, stabilization and mobilization of these. e) Various gene deletions causing defects in protein glycosylation or ER-associated protein degradation (ERAD) lead to an ER stress response, which has been associated with an increased number of LDs in the cell as well as an increase in TAG and SE content.

The family of perilipins, which are characterized by the presence of a conserved PAT (<u>P</u>erilipin/<u>A</u>DRP/<u>T</u>IP47) domain (Bickel, Tansey and Welte 2009), are usually one of the protein families specifically associated with the LD in those organisms that can accumulate high levels of neutral lipids (Ding *et al.* 2012; Zhu *et al.* 2015). As described above, perilipins can be associated with LD biogenesis, but can also have many other functions involved in stabilization and mobilization of the LD (Sztalryd and Brasaemle 2017). In yeasts and other fungal species, lipid droplet-associated perilipin-like proteins have also been recently identified, and while these are associated with lipid accumulating properties, their specific function remains still largely unknown (Wang and St Leger 2007; Zhu *et al.* 2015). A previously described LD-associated protein in *S. cerevisiae* (Pet10) was also recently identified as a perilipin and was shown to have an important role both at the level of LD assembly and post-assembly LD stabilization (Gao *et al.* 2017).

In this study, we propose that when *S. cerevisiae* cells reach high levels of TAG accumulation, there needs to be a simultaneous promotion of the cellular mechanisms of LD formation in the ER and/or further stabilization of this LD structure in order to unlock the further increase in TAG accumulation. To evaluate this hypothesis, we conducted this study on previously generated strains with highly increased metabolic fluxes towards TAG formation, able to accumulate up to 22% of their dry biomass as TAGs (Ferreira et al., 2018). Here, those strains were further engineered by focusing on targets involved with LD assembly and stabilization, such 3 as perilipins, perilipin-like fungal homologues, endogenous Fit2 homologues and genes that induce ER stress when deleted.

## 2 Results and Discussion

### 2.1 Improved TAG accumulation through expression of human and fungal Perilipins

In this study, we focused on further improvement of *S. cerevisiae* strains previously developed in our group able to accumulate levels of TAG up to 26-fold compared to a wild-type strain (Ferreira et al, 2018). These strains provide a platform with high metabolic flux towards TAG formation, therefore enabling us to investigate if LD assembly mechanisms can be a limiting factor in creating strains capable of accumulating even higher levels of TAG. The strain ADP was previously constructed by overexpressing either native or deregulated forms of enzymes known to be limiting steps in the metabolic pathways for fatty acid and TAG biosynthesis. To generate this strain, a genomic cassette was inserted expressing a deregulated mutant version of the acyl-CoA carboxylase gene *ACC1*, *ACC1^S659A/S1157A^* (*ACC1\*\**) (Shi *et al.* 2014) under control of the *HXT7* promoter, which is strongly expressed in low glucose (Teixeira *et al.* 2017). This strain also expresses an additional copy of the genes coding for phosphatidate phosphatase (*PAH1*) and the diacylglycerol acyltransferase (*DGA1*), expressed under control of the strong constitutive promoters P*_PGK1_* and P*_TEF1_*, respectively. This resulted in a strain reported previously to accumulate up to 13% of its dry biomass as TAGs. This strain was used to study the initial approaches suggested in this work since it has a high flux towards TAG production without resorting to a heavy engineering process and a fundamentally changed metabolic network. Furthermore, by using a strain that still contains all mechanisms involved in TAG hydrolysis, it is possible to detect targets that act on the same TAG degradation processes.

Expression of human perilipins in *S. cerevisiae* has been shown before to promote LD assembly, higher TAG synthesis fluxes and preventing their degradation by lipases (Jacquier *et al.* 2013). As a first step in our study, we investigated the effect that these same human perilipins would have on TAG levels when expressed in the ADP strain.

Perilipin-like proteins have also been identified in yeast and other fungi although not characterized to the same extent as they are in mammals and other animal species. To our knowledge, these encompass the *Rhodosporidium toruloides* Ldp1 (RtLdp1) (Zhu *et al.* 2015) and the *Metarhizium anisopliae* Mpl1 (MaMpl1) (Wang and St Leger 2007). These “perilipin-like” proteins have been shown to be LD-associated and correlated to neutral lipid production through loss-of-function studies, but their functions still remain mostly unknown. A *S. cerevisiae* perilipin Pet10 has also been recently described as an important target for LD assembly and structural stabilization during and after LD maturation (Gao *et al.* 2017).

To gain some insight into the function of these fungal perilipin-like proteins and simultaneously search for candidates better adapted to improving TAG accumulation in a yeast system, we performed genomic mining through BLASTP (http://blast.ncbi.nlm.nih.gov) using the RtLdp1 protein sequence as an input to search for homologues. From this search, 6 different candidates with arbitrarily different phylogenetic distance were picked for *in vivo* evaluation. The 5 new 6 candidate genes were termed: *MlPLP1* (Gene ID: 18931598) from *Melampsora laricipopulina* 98AG31, *LcPLP1* (GenBank: ORY92508.1) from *Leucosporidium creatinivorum*, *KbPLP1* (Gene ID: 27417221) from *Kalmanozyma brasiliensis* GHG001, *XdPLP1* (GenBank: CED83179.1) from *Xanthophyllomyces dendrorhous*, *OrPLP1* (GenBank: OCH92582.1) from *Obba rivulosa*, *MoPLP1* (GenBank: GAA98449.1) from *Mixia osmundae* IAM 14324.

The 3 human perilipin genes *PLIN1*, *PLIN2/ADRP* and *PLIN3/TIP47*, the 6 newly identified candidates, the 3 previously described fungal perilipins *RtLDP1*, *MaMPL1*, *PET10* and a previously described putative perilipin-like protein YALI0F24167p from *Yarrowia lipolytica* (Gene ID: 2908726), here named *YlPLP1*, were obtained as synthetic genes with codon-optimized sequences for expression in *S. cerevisiae*. These genes were individually cloned in a centromeric single copy plasmid (p416) with the strong constitutive promoter P*_TEF1_*. Plasmids were used to transform the ADP strain using the empty plasmid p416TEF as a control.

TAG levels were analyzed for each resulting strain after cultivation in minimal medium with 2% glucose for 72 h in order to evaluate the effects on TAG production and storage capacity after glucose and ethanol were consumed and the cells allowed to enter stationary phase. In the resulting strains, final TAG levels were increased up to 58.7 mg/g DCW (dry cell weight) (Figure 2). Compared to the control strain with an empty plasmid, TAG levels were increased by 22%, 23% or 28% for *PLIN1, 2* or *3* respectively. From the 10 screened fungal perilipin homologue genes, only 3 of them showed an increase in accumulated TAGs when expressed in ADP: *MlPLP1*, *KbPLP1* and *MoPLP1* which increased final TAG accumulated levels by 25%, 26% and 14%, respectively. None of the previously described fungal perilipin genes *RtLDP1*, *MaMPL1* or *PET10* improved TAG production in this strain (Figure 2).

**Figure 2.**
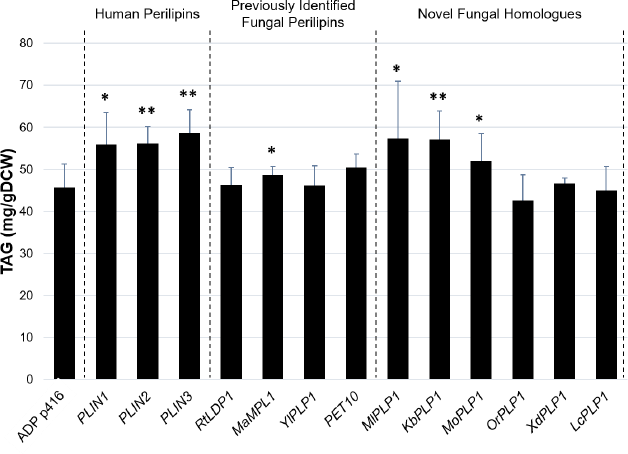
TAG production of *S. cerevisiae* strains expressing different perilipin and perilipin-related proteins. Codon-optimized versions of different perilipin homologues were expressed from a centromeric plasmid in a previously engineered strain ADP overexpressing *ACC1*\*\*, *PAH1* and *DGA1* for accumulation of high TAG levels. Three different categories are represented: Human perilipins: PLIN1, PLIN2/ADRP and PLIN3/TIP47; Fungal perilipin-like protein homologues identified in the literature: RtLDP1 from *Rhodosporidium toruloides*, MaMPL1 from *Metarhizium anisopliae*, YlPLP1 from *Yarrowia lipolytica* and Pet10 from *Saccharomyces cerevisiae*; and 6 novel candidates identified from a bioinformatic analysis using BLASTP: MlPLP1 from *Melampsora larici-populina* 98AG31, KbPLP1 from *Kalmanozyma brasiliensis*, MoPLP1 from *Mixia osmundae* IAM 14324, OrPLP1 from *Obba rivulosa*, XdPLP1 from *Xanthophyllomyces dendrorhous* and LcPLP1 from *Leucosporidium creatinivorum*. Resulting strains were cultivated in minimal medium with 2% glucose for 72 h. Plot shows average and error bars which represent standard deviation from at least 3 biological replicates. *p-value < 0.05, **p-value < 0.005 (Student’s t test, one-tailed, unequal variance)

These results suggest that perilipins provide beneficial effects either at the level of the LD assembly process, therefore becoming more efficient at storing produced TAGs, or at stabilizing the lipid droplet after it is formed, such as protecting it from lipases, autophagy or other possible degradation mechanisms (Kaushik and Cuervo 2015; Sztalryd and Brasaemle 2017). Furthermore, analysis of fungal perilipins seems to show some disparity between these. While some homologues are able to confer the same improvement in TAG accumulation capacity as human perilipins, most of them do not seem to have a significant effect. This could indicate a difference in function of these different proteins as it is seen for mammalian perilipins where different homologues can be involved in different processes concerning neutral lipid storage or mobilization (Kimmel and Sztalryd 2016).

Because an unexpected difference was seen between the TAG levels of the uracil-prototrophic strain ADP with plasmid p416 and the levels observed in our previous study where uracil-auxotrophic strains were grown with supplemented uracil in the medium, a control experiment was performed comparing simultaneous lipid extraction of both cultures. Analysis of TAG content after extraction showed that uracil-prototrophic strains (carrying the p416 plasmid) consistently produced up to 40% less TAGs than their uracil-fed auxotrophic counterparts (Supplementary Figure S1). Furthermore, we observed TAG levels of 81 mg/g DCW and 150 mg/g DCW for ADP and RF07, respectively, instead of the previously reported 129 mg/gDCW and 218 mg/gDCW respectively. Even though this was a very significant decrease, for the sake of a simpler study design, increased strain construction efficiency and time advantage, evaluation of gene expression using a plasmid system was kept as the main strategy.

### 2.2 Plin3 and MlPlp1 improve TAG accumulation even when lipases are absent

Some perilipins, such as the human Plin1, have been described to act in neutral lipid storage by blocking the access of lipases and other hydrolytic enzymes to the lipids in the lipid droplet(“shielding”) (Brasaemle *et al.* 2000; Sztalryd and Brasaemle 2017). In order to evaluate if the improvements in TAG levels are due to an effect of lipid droplet shielding from lipases, the two most beneficial fungal perilipin homologues *MlPLP1, KbPLP1,* the most beneficial human perilipin gene *PLIN3* and the native *S. cerevisiae* perilipin gene *PET10* were expressed in strain RF07. RF07 was derived from the ADP strain by deleting the main lipase genes *TGL3, 4* and *5*, which resulted in TAG levels up to 84% higher than for ADP (Supplementary Figure S1).

Expression of *PLIN3* in RF07 resulted in an improvement of 34% in the final TAG levels over RF07 with an empty plasmid. This increase in percentage was an even higher increase than when this gene was expressed in ADP. *MlPLP1* expression increased TAG levels by 22%, a value comparable to its expression in ADP, while *KbPLP1* expression only increased TAG content by 9% in a statistically non-significant way (p value > 0.05). As in ADP, overexpression of *PET10* did not evoke a beneficial effect on TAG accumulation (Figure 3 A).

**Figure 3.**
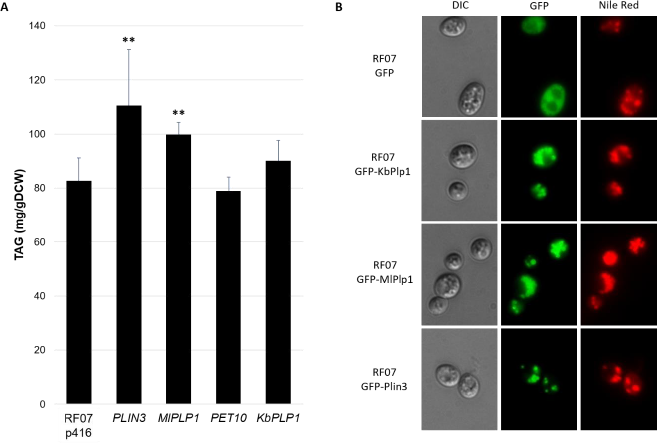
Effect of perilipin expression in a high TAG accumulating strain lacking lipases. A) Codon-optimized versions of different perilipin homologues were expressed in RF07 (ADP Δ*tgl3/4/5*). Resulting strains were cultivated in minimal medium with 2% glucose for 72 h. Plot shows average and error bars which represent standard deviation from at least 4 biological replicates. *p-value < 0.05, **p-value < 0.005 (Student’s t test, one-tailed, unequal variance). b) Fluorescence microscopy of RF07 cells expressing GFP-tagged perilipins and stained with Nile Red after 60 h cultivation in minimal medium with 2% glucose.

These results suggest firstly that *PLIN3* and *MlPLP1* probably act on improving TAG accumulation not by shielding the lipid droplet from lipase hydrolysis, but rather by LD assembly and/or budding. This is consistent with previous reports for Plin3 in which this protein is described to be natively present in nascent lipid droplets and involved in lipid droplet generation rather than involved in shielding LDs from lipases (Wolins *et al.* 2005; Bulankina *et al.* 2009; Sztalryd and Brasaemle 2017). On the other hand, *KbPLP1* expression has a decreased effect when the main lipases are not present compared to when it is expressed in ADP, indicating that its primary benefit in TAG accumulation might rely on a capacity to shield the lipid droplet from enzymes with lipase activity. The 9% improvement in *KbPLP1* might derive either from a slight improvement in LD biogenesis aspects or protection from other enzymes with less prominent lipase activity.

Localization of these perilipin proteins was analyzed by expressing a N-terminally GFP fusion to these proteins. The same cells were stained with Nile Red, a dye specific for staining of lipids with high affinity for neutral lipids such as TAGs. With Nile Red staining, it is possible to identify lipid droplets due to their high TAG content. Analysis of these strains with fluorescent microscopy shows that GFP-tagged perilipins appear as distinct foci overlapping with the position of stained lipid droplets, confirming that these do localize to lipid droplets when expressed in RF07 (Figure 3 B). This is congruent with previous reports of Plin3 localization when expressed in *S. cerevisiae* (Jacquier *et al.* 2013).

### 2.3 Absence of seipin Sei1 allows for increased accumulation of TAGs

The seipin Sei1 (often referred to as Fld1) has been shown in *S. cerevisiae* to be involved in releasing formed lipid droplets and detaching these from the ER (Wolinski *et al.* 2015). Deletion of this gene has been associated with formation of abnormal LD structures: either “supersized lipid droplets” with a single or low amount of lipid droplets with large volumes instead of several small ones (Szymanski *et al.* 2007; Fei *et al.* 2011; Wolinski *et al.* 2011), or clusters of small irregular droplets embedded in locally proliferated ER membranes (Wolinski *et al.* 2011; Cartwright *et al.* 2015; Grippa *et al.* 2015). The *Δsei1* phenotype has also been associated with increased TAG and SE content (Fei *et al.* 2008). Lipid droplet assembly, budding and release is a dynamic process and all its components need to be timed properly in order to optimize TAG formation fluxes and accumulation of the product in LDs.

We evaluated how TAG accumulation is changed when segregation of LDs from the ER is severely defective. This was implemented by deleting the seipin gene *SEI1* from RF07.

Deletion of *SEI1* led to an increase in TAG accumulation to 141 mg/gDCW, a 64% increase over RF07 (Figure 4 A). This suggests that for optimal TAG accumulation, either formation of large lipid droplets is favourable, or there is a benefit from the LDs being associated with the ER for a longer time. A potential benefit of increased association time with the ER might be that during this time there is a conversion of PA to DAG and consequently TAG that is transferred to the attached LD. Therefore, a longer attachment time might result in a prolonged steady flow of TAGs towards the LD, enabling further accumulation.

**Figure 4.**
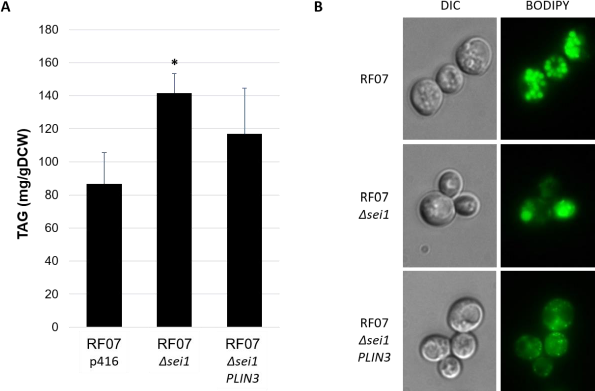
Effects deleting the seipin gene *SEI1* in strain RF07. A) Impact of *SEI1* deletion and subsequent expression of *PLIN3* on TAG accumulation in RF07. Strains were cultivated in minimal medium with 2% glucose for 72 h. Plot shows average and error bars which represent standard deviation from at least 4 biological replicates. *p-value < 0.05 (Student’s t test, one-tailed, unequal variance). B) Fluorescent microscopy of Δ*sei1* strains. Cells were stained with BODIPY after 60 h cultivation in minimal medium with 2% glucose.

In order to address the combinatorial effect of promoting LD budding through perilipin expression and extended association with the ER, *PLIN3* was expressed in this same strain. However, TAG levels were decreased in comparison with RF07 *Δsei1*.

When looking at the lipid droplet structure through fluorescence microscopy, the structure of these was altered between the three different strains (Figure 4 B). Deletion of *SEI1* in RF07 creates the previously observed phenotype where lipid droplets have an abnormal physiology of either a single supersized lipid droplet or a cluster of lipid droplets that spread a diffuse fluorescence signal. Curiously, microscopy observation of *Δsei1 PLIN3* mutants shows many cells with small intracellular disperse lipid droplets stained by BODIPY and a diffuse fluorescence across the cytoplasm. The results may indicate an interference from Plin3 in lipid droplet formation when Sei1 is absent, correlating also with amorphous morphology of the ER-LD contact dynamics. To our knowledge, no studies have reported this phenotype or seipin-perilipin interactions that could lead to this phenomenon.

### 2.4 Overexpression of the FIT2 homologue *YFT2* improves TAG accumulation

Fat storage-Inducing Transmembrane (FIT) proteins have been shown to represent major factors for budding of the ER membrane during LD biogenesis and absence of these has been shown to create droplets inside the ER membrane without proper budding (Choudhary *et al.* 2015). Supporting this, when the mouse FIT2 protein was expressed in plants, it was reported to induce high lipid droplet accumulation (Cai *et al.* 2017).

Since Plin3 has been shown to be involved in promoting LD budding and expression of the protein resulted in improved TAG production in our strains, we hypothesized that LD budding kinetics might be a limiting factor for TAG accumulation in strains with high metabolic fluxes towards TAG production. In order to evaluate this, we proposed that overexpression of the *S. cerevisiae* FIT2 homologues *SCS3* and *YFT2* could improve TAG accumulation through promotion of LD budding.

*SCS3* and *YFT2* were expressed from the plasmid p416TEF in RF07. Overexpression of *YFT2* improved TAG accumulation levels by 27% to 104 mg/gDCW, while expression of *SCS3* did not result in a significant difference in TAG levels (Figure 5 A).

**Figure 5.**
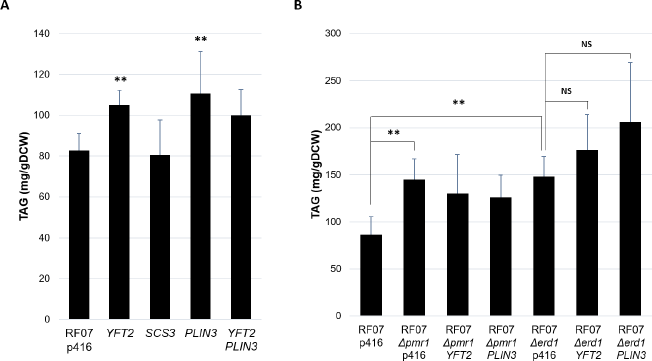
Effect of FIT protein overexpression and promoting ER stress on TAG accumulation. A) Endogenous FIT protein genes *YFT2* or *SCS3*, and/or human perilipin gene *PLIN3* were expressed in RF07 (ADP Δ*tgl3/4/5*). B) Genes *ERD1* and *PMR1* involved in ER stress were deleted. Additional expression of *YFT2* or *PLIN3* was also evaluated in these deletion strains. Resulting strains were cultivated in minimal medium with 2% glucose for 72 h. Plots show average and error bars which represent standard deviation from at least 4 biological replicates. *p-value < 0.05, **p-value < 0.005, NS: not statistically significant (Student’s t test, one-tailed, unequal variance).

Although Scs3 and Yft2 are homologues (Choudhary *et al.* 2015), the two proteins have been described as functionally different when knockout mutants were analyzed through transcriptomics (Moir *et al.* 2012). The results from *YFT2* and *SCS3* expression in RF07 further support the existence of these functional differences between the two genes.

In order to access if there is an additive or synergistic effect between *PLIN3* and *YFT2*, we cloned both genes on the same p416 plasmid under control of strong constitutive promoters (P_TEF1_ and P_PGK1_) and the TAG levels in resulting strains were analyzed as before. Simultaneous expression of the two genes improved TAG accumulation by 24%, close to the same levels as expressing only *YFT2*, and a lower level than expressing *PLIN3* alone.

Since Yft2 is specifically involved in LD budding and assembly, we assume that LD budding is indeed a mechanism that can be limiting TAG accumulation in our strain, and overexpression of *YFT2* might overcome this limitation. Furthermore, since PLIN3 has been shown to act mostly on the LD assembly process in mammals, we think that it might play the same role when expressed in our strains. We therefore speculate that improved LD assembly and budding might be the main cause of the observed Plin3-mediated improvement in RF07. This is supported by the fact that simultaneous expression of *YFT2* and *PLIN3* does not further improve TAG titers. Assuming that both are acting on the same overall process even if through different mechanisms, we consider it a fair assumption that expression of only one of them would be sufficient to relieve the limiting step in LD assembly.

### 2.5 TAG accumulation is further improved by promoting endoplasmic reticulum stress

The data obtained in this study suggest that improving or further stimulating lipid droplet formation mechanisms led to increased TAG accumulation in our strains which have high fluxes towards TAG biosynthesis.

Conditions of ER stress have been strongly associated with improved LD formation and increased TAG content in *S. cerevisiae* (Fei *et al.* 2009; Moir *et al.* 2012). Various gene deletions causing defects in protein glycosylation or ER-associated protein degradation lead to an ER stress response and have been associated with an increased number of LDs in the cell as well as an increase in TAG and SE content (Uchimura, Sugiyama and Nikawa 2005; Fei *et al.* 2009; Chantret *et al.* 2011). Using this knowledge, we individually deleted in strain RF07 two of these identified genes, *ERD1* encoding for a predicted membrane protein required for lumenal ER protein retention, and *PMR1* encoding for a Ca^2+^/Mn^2+^ P-type ATPase. The deletion phenotypes of these genes were both previously associated with an increase in TAG content and increased LD number (Fei *et al.* 2009) while tagged as “viable” on SGD (*Saccharomyces Genome Database*, https://www.yeastgenome.org). In order to evaluate if these processes are additive or epistatic with the previously identified phenotypes, the gene deletions were evaluated also in combination with overexpression of either *YFT2* or *PLIN3*.

Analysis of the newly generated strains showed TAG levels of 148 mg/gDCW and 145 mg/gDCW, an increase of 72% and 67% in TAG content when *ERD1* or *PMR1*, respectively, were deleted in RF07. When additionally overexpressing *YFT2* in the Δ*erd1* strain, the TAG content was increased by 104% compared to RF07 and when *PLIN3* was expressed the increase over RF07 was of 138%, reaching an average level of 201 mg/gDCW (Figure 5 B). However, due to high variation of the obtained data points for these mutants, the effect of overexpression of both *YFT2* and *PLIN3* was not statistically significant when compared to RF07 *Δerd1*, although it is if compared to RF07. Expression of either *YFT2* or *PLIN3* in the *Δpmr1* strain did not result in an 13 increase in TAG content. Furthermore, measurements of OD_600_ showed that deletion of *ERD1* only had a minor impact on final cell density, showing OD values of 10.8 for *Δerd1* and 13.2 for RF07, while deletion of *PMR1* had a drastic effect on final cell density, reducing the OD to 5.4, a 60% decrease over RF07 (Supplementary Figure S2). In fact, decrease of growth rate due to deletion of *PMR1* has been reported in literature since this causes numerous defects in Ca^2+^ dependent processes (Antebi and Fink 1992).

Observation of lipid droplet morphology on the strains RF07 *Δerd1 and RF07 Δpmr1* using BODIPY fluorescence staining and fluorescence microscopy did not reveal any evident phenotypic changes compared to RF07 (data not shown).

The increase in TAG content observed after deletion of *ERD1* or *PMR1* was similar to the one observed earlier (Fei *et al.* 2009). However, it is important to note that in the mentioned study these mutations were analysed in a wild-type strain, where the TAG content in the cell is less than 7 % of the TAG levels in RF07 (Ferreira et al., 2018).

These results indicate once again the importance of the mechanisms involved in LD formation from the ER in enabling further TAG accumulation in yeast. The two gene deletions applied have been shown to cause ER stress and through this stimulate LD formation. The relationship between these two processes is thought to be because the increase in LDs surface might provide an improved capacity to accommodate larger amounts of unfolded proteins which have exposed hydrophobic parts, this way preventing them from aggregation and thus protecting cell viability (Fei *et al.* 2009). This process has been shown to be independent from the action of Ire1 and consequently the unfolded protein response (UPR). Furthermore, it also did not show higher protein levels for Dga1, Lro1, Are1 and Are2 in the cell extracts, indicating an independence of upregulating any storage lipid synthesis (Patil and Walter 2001). As such, the signaling mechanisms between the two processes: ER stress and LD budding, remain to be elucidated.

## 3 Conclusions

In this work we successfully engineered yeast strains for high levels of TAG production and accumulation through targeting of processes involved in lipid droplet biogenesis and assembly. For that we (over)expressed and deleted genes involved in physiological and structural mechanisms of lipid droplet assembly and budding such as perilipins, FIT proteins and proteins involved in stimulating ER stress.

The engineering strategy employed in this work differs from common metabolic engineering approaches where the enzymes of a pathway are the main focus of attention. Here, we instead targeted mostly structural aspects of the organelles where the enzymes involved in the pathway are localized. The work resulted in strains with 35-fold increase in TAG production capable of producing up to 21% of their dry biomass as TAGs, highlighting the importance of considering the whole cellular context of lipid droplets when engineering pathways that heavily depend or interact with this organelle.

Lipid droplets are a complex structure with many undescribed mechanisms regulating their dynamics. A deep understanding of this organelle as well as its biogenesis, degradation and interactions is crucial as starting point for engineering of lipid metabolism processes in the development of yeast as a cell factory. This was here shown to be true for production of TAGs, as it might be for many lipid and fatty acid-derived products. Hydrophobic surfaces such as membranes and LDs are the most probable intracellular site where hydrophobic products and intermediaries will be stored when being overproduced. Expansion of cellular membrane surface was previously explored as a viable strategy to increase carotenoid production in E. coli (Wu *et al.* 2017). In the same way, we envision that expansion of the LD volume and number in the cell can promote further production of hydrophobic compounds by providing an intracellular storage facility that is segregated from membranar cell functions.

Through this study we also generated valuable insight concerning the function of the different components used. The generated data on the tested fungal perilipins provides new knowledge about a mostly undescribed set of proteins, providing insight on the possible action of these and their benefits towards lipid accumulation in the native hosts. Furthermore, the role of Yft2 and Scs3 and their relationship to the ER stress phenotypes generated are of utmost interest due to their conserved character among eukaryotes. The results generated here not only show the relevance of Yft2 for LD assembly and TAG biosynthesis but also contribute towards understanding its differences to Scs3 and the mechanism of action for both these proteins in the LD assembly process.

In conclusion, this work is a successful approach towards development of yeast as a cell factory for TAG production. Combination of the factors applied here with previous approaches for increasing flux towards the TAG biosynthetic pathway brings *S. cerevisiae* closer to be an industrially viable organism for this process as it is able to compete with the efficiency of other leading organisms in this front. Furthermore, it is possible to combine these efforts with changes in fatty acid machinery and in specificity of the enzymes involved in TAG biosynthesis to enable production of tailored TAGs with specially desired properties and increased value.

## 4 Materials and Methods

### 4.1 Plasmid and strain construction

Genes *PLIN1*, *PLIN2*, *PLIN3*, *RtLDP1*, *MaMPL1* and *YlPLP1* were ordered as codon-optimized synthetic genes and *MlPLP1*, *KbPLP1*, *MoPLP1*, *OrPLP1*, *XdPLP1* and *LcPLP1* as codon-optimized GenParts from Genscript (Piscataway, NJ, USA). Genes *PET10*, *YFT2* and *SCS3* were amplified from *S. cerevisiae* CEN.PK113-5D genomic DNA.

Genes were amplified by PCR to generate the complementary overhangs for insertion into the plasmid by Gibson assembly (New England Biolabs, Ipswich, MA, USA) and cloned into p416TEF (ATCC^®^ 87368™) linearized by the restriction enzymes *Bam*HI and *Xho*I. For simultaneous cloning of *YFT2* and *PLIN3* all promoter, terminator and gene fragments were amplified by PCR, fused by fusion PCR and then cloned into the respective plasmid by Gibson assembly. GenParts did not require a PCR amplification and were cloned directly using Gibson assembly in the same way.

Yeast transformations were performed using lithium acetate and PEG3350 as described before (Gietz and Schiestl 2007). A list of strains, primers used for plasmid construction, generated plasmids, synthetic gene and GenParts sequences are shown in **Additional File 1**.

### 4.2 Growth medium

ADP and RF07 strains were kept on YPD plates containing 20 g/L glucose, 10 g/L yeast extract, 20 g/L peptone from casein and 20 g/L agar. Plasmid-carrying strains were grown on selective growth medium containing 6.9 g/L yeast nitrogen base w/o amino acids (Formedium, Hunstanton, UK), 0.77 g/L complete supplement mixture w/o uracil (Formedium), 20 g/L glucose and 20 g/L agar.

Shake flask cultivations were performed in minimal medium containing 20 g/L glucose, 5 g/L (NH_4_)_2_SO_4_, 14.4 g/L KH_2_PO_4_, 0.5 g/L MgSO_4_∙7H_2_O. After sterilization, 2 mL/L trace element solution and 1mL/L of vitamin solution were added. The composition of the trace element and vitamin solution has been reported earlier (Verduyn *et al.* 1992). When growing strains with a uracil auxotrophy, 60 mg/L of uracil was added to the medium.

### 4.3 Shake flask cultivations

All experiments were performed with strains cultivated as 4 biological replicates, i.e. 4 independent transformants were used to start pre-cultures. Some of the experiments were repeated up to 3 times for increasing statistical power of high variating strains. For these, 3 mL of minimal medium in a 15 mL tube were inoculated and cultivated at 200 rpm and 30°C for 24-40 h. Subsequently, the pre-culture was used to inoculate 15 mL of minimal medium in a 100 mL shake flask at a starting OD_600_ of 0.1. Shake flasks were incubated at 200 rpm and 30°C for up to 72 h.

A spectrophotometer (Genesis20, Thermo Fisher Scientific, Waltham, MA, USA) was used to measure cell growth at designated time points and at the end of the shake flask cultivations. Optical density (OD_600_) was measured by absorbance at 600nm of a diluted culture sample.

### 4.4 Lipid Quantification

Samples for lipid analysis were taken as 10 mL of culture at the end of the shake flask cultivations, after 72 h. Subsequently, the samples were centrifuged at 4000 rpm and the supernatant was discarded. The pellets were kept at −80°C for 10 min and then freeze-dried using a Christ alpha 2-4 LSC (Christ Gefriertrocknungsanlagen, Osterode, Germany). The samples were extracted using a microwave-assisted CHCl_3_:MeOH method and analyzed by HPLC-CAD as described previously (Khoomrung *et al.* 2013). For lipid extraction, 5 to 10 mg of dry cell biomass was used with 200 μg of cholesterol as internal standard. Quantification of lipids was performed using Chromeleon software (Thermo Fisher Scientific, Waltham, MA, USA) and further normalization to both internal standard and biomass values.

### 4.5 Neutral Lipid Staining

100 μL of cell culture were transferred into a 1.5 mL tube, centrifuged and the pellet washed with 1 mL of deionized water. Cells were then centrifuged at 3000 g for 5 minutes and resuspended in 100 μL phosphate-buffered saline (PBS). Resuspended cells were treated with 1 μL of BODIPY 493/503 (Thermo Fisher Scientific, Waltham, MA, USA) solution (1 mg/mL in ethanol) or 3 μL Nile Red (1 mg/mL in DMSO) and kept at 4°C in the dark for 30 minutes. Fluorescent microscope pictures were taken using a Leica DMI4000 B inverted microscope (Leica Microsystems) and processed with the Leica Application Suite (LAS AF 6000 E) software.

## Acknowledgments

We thank the Chalmers Mass Spectrometry Infrastructure team for their help with the HPLC-CAD analysis. We also thank Dr. Michael Gossing and David Bergenholm for helpful feedback during this project.

## Contributions

P.G.T, V.S, F.D and J.N conceived and designed the project. P.G.T performed the experiments. The manuscript was written by P.G.T. and edited by V.S, F.D and J.N. All authors read and approved the manuscript.

## Supplementary Figures

**Supplementary Figure S1.**
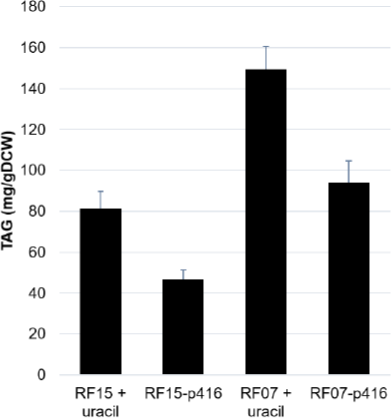
Effects in TAG accumulation of uracil-auxotrophic strains grown in minimal media versus the same strain with an empty centromeric plasmid. Strains were cultivated in minimal medium with 2% glucose for 72 h. Plot shows average and error bars which represent standard deviation from at least 3 biological replicates.

**Supplementary Figure S2.**
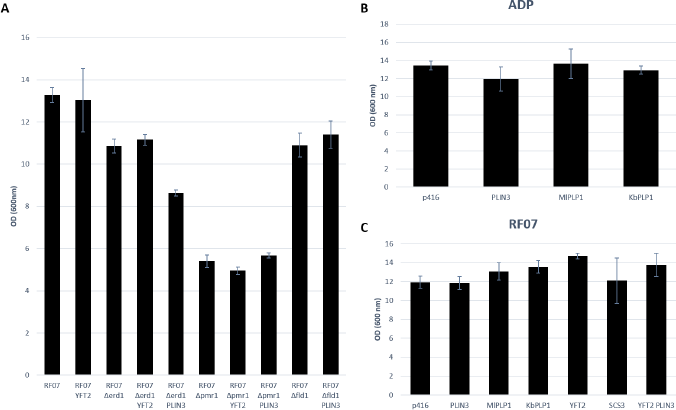
Final OD_600_ values of strains used during this study. Strains were cultivated in minimal medium with 2% glucose for 72 h. Plot shows average and error bars which represent standard deviation from at least 3 biological replicates. 24

## References

Adeyo O, Horn PJ, Lee S et al. The yeast lipin orthologue Pah1p is important for biogenesis of lipid droplets. J Cell Biol 2011;192:1043–55.

Antebi A, Fink GR. The yeast Ca(2+)-ATPase homologue, PMR1, is required for normal Golgi function and localizes in a novel Golgi-like distribution. Mol Biol Cell 1992;3:633–54.

Barbosa AD, Sembongi H, Su W-M et al. Lipid partitioning at the nuclear envelope controls membrane biogenesis. Mol Biol Cell 2015;26:3641–57.

Beller M, Thiel K, Thul PJ et al. Lipid droplets: a dynamic organelle moves into focus. FEBS Lett 2010;584:2176–82.

Bickel PE, Tansey JT, Welte MA. PAT proteins, an ancient family of lipid droplet proteins that regulate cellular lipid stores. Biochim Biophys Acta 2009;1791:419–40.

Brasaemle DL, Rubin B, Harten IA et al. Perilipin A increases triacylglycerol storage by decreasing the rate of triacylglycerol hydrolysis. J Biol Chem 2000;275:38486–93.

Brasaemle DL, Wolins NE. Packaging of fat: an evolving model of lipid droplet assembly and expansion. J Biol Chem 2012;287:2273–9.

Bulankina AV, Deggerich A, Wenzel D et al. TIP47 functions in the biogenesis of lipid droplets. J Cell Biol 2009;185:641–55.

Cai Y, McClinchie E, Price A et al. Mouse fat storage-inducing transmembrane protein 2 (FIT2) promotes lipid droplet accumulation in plants. Plant Biotechnol J 2017;15:824–36.

Cartwright BR, Binns DD, Hilton CL et al. Seipin performs dissectible functions in promoting lipid droplet biogenesis and regulating droplet morphology. Mol Biol Cell 2015;26:726–39.

Chantret I, Kodali VP, Lahmouich C et al. Endoplasmic reticulum-associated degradation (ERAD) and free oligosaccharide generation in *Saccharomyces cerevisiae*. J Biol Chem 2011;286:41786–800.

Chen X, Goodman JM. The collaborative work of droplet assembly. Biochim Biophys Acta 2017;1862:1205–11.

Choudhary V, Ojha N, Golden A et al. A conserved family of proteins facilitates nascent lipid droplet budding from the ER. J Cell Biol 2015;211:261–71.

Czabany T, Athenstaedt K, Daum G. Synthesis, storage and degradation of neutral lipids in yeast. Biochim Biophys Acta 2007;1771:299–309.

Ding Y, Wu Y, Zeng R et al. Proteomic profiling of lipid droplet-associated proteins in primary adipocytes of normal and obese mouse. Acta Biochim Biophys Sin 2012;44:394–406.

d’Espaux L, Ghosh A, Runguphan W et al. Engineering high-level production of fatty alcohols by *Saccharomyces cerevisiae* from lignocellulosic feedstocks. Metab Eng 2017;42:115–25.

Fei W, Shui G, Gaeta B et al. Fld1p, a functional homologue of human seipin, regulates the size of lipid droplets in yeast. J Cell Biol 2008;180:473–82.

Fei W, Shui G, Zhang Y et al. A role for phosphatidic acid in the formation of “supersized” lipid droplets. PLoS Genet 2011;7:e1002201.

Fei W, Wang H, Fu X et al. Conditions of endoplasmic reticulum stress stimulate lipid droplet formation in *Saccharomyces cerevisiae*. Biochem J 2009;424:61–7.

Fernandez-Moya R, Da Silva NA. Engineering *Saccharomyces cerevisiae* for high-level synthesis of fatty acids and derived products. FEMS Yeast Res 2017;17, DOI: 10.1093/femsyr/fox071.

Ferreira R, Teixeira PG, Gossing M et al. Metabolic engineering of Saccharomyces cerevisiae for overproduction of triacylglycerols. Metabolic Engineering Communications 2018/6;6:22–7.

Galafassi S, Cucchetti D, Pizza F et al. Lipid production for second generation biodiesel by the oleaginous yeast *Rhodotorula graminis*. Bioresour Technol 2012;111:398–403.

Gao Q, Binns DD, Kinch LN et al. Pet10p is a yeast perilipin that stabilizes lipid droplets and promotes their assembly. J Cell Biol 2017, DOI: 10.1083/jcb.201610013.

Gietz RD, Schiestl RH. High-efficiency yeast transformation using the LiAc/SS carrier DNA/PEG method. Nat Protoc 2007;2:31–4.

Grippa A, Buxó L, Mora G et al. The seipin complex Fld1/Ldb16 stabilizes ER-lipid droplet contact sites. J Cell Biol 2015;211:829–44.

Han G-S, Wu W-I, Carman GM. The *Saccharomyces cerevisiae* Lipin homolog is a Mg2+-dependent phosphatidate phosphatase enzyme. J Biol Chem 2006;281:9210–8.

Jacquier N, Mishra S, Choudhary V et al. Expression of oleosin and perilipins in yeast promotes formation of lipid droplets from the endoplasmic reticulum. J Cell Sci 2013;126:5198–209.

Joshi AS, Zhang H, Prinz WA. Organelle biogenesis in the endoplasmic reticulum. Nat Cell Biol 2017;19:876–82.

Karanasios E, Han G-S, Xu Z et al. A phosphorylation-regulated amphipathic helix controls the membrane translocation and function of the yeast phosphatidate phosphatase. Proc Natl Acad Sci U S A 2010;107:17539–44.

Kaushik S, Cuervo AM. Degradation of lipid droplet-associated proteins by chaperone-mediated autophagy facilitates lipolysis. Nat Cell Biol 2015;17:759–70.

Khoomrung S, Chumnanpuen P, Jansa-Ard S et al. Rapid quantification of yeast lipid using microwave-assisted total lipid extraction and HPLC-CAD. Anal Chem 2013;85:4912–9.

Kimmel AR, Sztalryd C. The Perilipins: Major Cytosolic Lipid Droplet-Associated Proteins and Their Roles in Cellular Lipid Storage, Mobilization, and Systemic Homeostasis. Annu Rev Nutr 2016;36:471–509.

Kohlwein SD, Veenhuis M, van der Klei IJ. Lipid Droplets and Peroxisomes: Key Players in Cellular Lipid Homeostasis or A Matter of Fat-Store ‘em Up or Burn ‘em Down. Genetics 2013;193:1–50.

Krivoruchko A, Nielsen J. Production of natural products through metabolic engineering of *Saccharomyces cerevisiae*. Curr Opin Biotechnol 2015;35:7–15.

Moir RD, Gross DA, Silver DL et al. SCS3 and YFT2 link transcription of phospholipid biosynthetic genes to ER stress and the UPR. PLoS Genet 2012;8:e1002890.

Natter K, Kohlwein SD. Yeast and cancer cells - common principles in lipid metabolism. Biochim Biophys Acta 2012;1831:314–26.

Nielsen J, Keasling JD. Engineering Cellular Metabolism. Cell 2016;164:1185–97.

Patil C, Walter P. Intracellular signaling from the endoplasmic reticulum to the nucleus: the unfolded protein response in yeast and mammals. Curr Opin Cell Biol 2001;13:349–55.

Petranovic D, Tyo K, Vemuri GN et al. Prospects of yeast systems biology for human health: integrating lipid, protein and energy metabolism. FEMS Yeast Res 2010;10:1046–59.

Pfleger BF, Gossing M, Nielsen J. Metabolic engineering strategies for microbial synthesis of oleochemicals. Metab Eng 2015;29:1–11.

Santos‐ Rosa H, Leung J, Grimsey N et al. The yeast lipin Smp2 couples phospholipid biosynthesis to nuclear membrane growth. EMBO J 2005;24:1931–41.

Schuldiner M, Bohnert M. A different kind of love - lipid droplet contact sites. Biochim Biophys Acta 2017;1862:1188–96.

Shi S, Chen Y, Siewers V et al. Improving production of malonyl coenzyme A-derived metabolites by abolishing Snf1-dependent regulation of Acc1. MBio 2014;5:e01130–14.

Sztalryd C, Brasaemle DL. The perilipin family of lipid droplet proteins: Gatekeepers of intracellular lipolysis. Biochim Biophys Acta 2017;1862:1221–32.

Szymanski KM, Binns D, Bartz R et al. The lipodystrophy protein seipin is found at endoplasmic reticulum lipid droplet junctions and is important for droplet morphology. Proc Natl Acad Sci U S A 2007;104:20890–5.

Teixeira PG, Ferreira R, Zhou YJ et al. Dynamic regulation of fatty acid pools for improved production of fatty alcohols in *Saccharomyces cerevisiae*. Microb Cell Fact 2017;16:45.

Thiam AR, Beller M. The why, when and how of lipid droplet diversity. J Cell Sci 2017;130:315–24.

Tsigie YA, Wang C-Y, Truong C-T et al. Lipid production from *Yarrowia lipolytica* Po1g grown in sugarcane bagasse hydrolysate. Bioresour Technol 2011;102:9216–22.

Uchimura S, Sugiyama M, Nikawa J-I. Effects of N-glycosylation and inositol on the ER stress response in yeast *Saccharomyces cerevisiae*. Biosci Biotechnol Biochem 2005;69:1274–80.

Verduyn C, Postma E, Scheffers WA et al. Effect of benzoic acid on metabolic fluxes in yeasts: a continuous-culture study on the regulation of respiration and alcoholic fermentation. Yeast 1992;8:501–17.

Wang C, St Leger RJ. The *Metarhizium anisopliae* Perilipin Homolog MPL1 Regulates Lipid Metabolism, Appressorial Turgor Pressure, and Virulence. J Biol Chem 2007;282:21110–5.

Welte MA. Expanding roles for lipid droplets. Curr Biol 2015;25:R470–81.

Wilfling F, Haas JT, Walther TC et al. Lipid droplet biogenesis. Curr Opin Cell Biol 2014;29:39–45.

Wilfling F, Wang H, Haas JT et al. Triacylglycerol synthesis enzymes mediate lipid droplet growth by relocalizing from the ER to lipid droplets. Dev Cell 2013;24:384–99.

Wolinski H, Hofbauer HF, Hellauer K et al. Seipin is involved in the regulation of phosphatidic acid metabolism at a subdomain of the nuclear envelope in yeast. Biochim Biophys Acta 2015;1851:1450–64.

Wolinski H, Kolb D, Hermann S et al. A role for seipin in lipid droplet dynamics and inheritance in yeast. J Cell Sci 2011;124:3894–904.

Wolins NE, Quaynor BK, Skinner JR et al. S3-12, Adipophilin, and TIP47 package lipid in adipocytes. J Biol Chem 2005;280:19146–55.

Wu T, Ye L, Zhao D et al. Membrane engineering - A novel strategy to enhance the production and accumulation of β-carotene in *Escherichia coli*. Metab Eng 2017;43:85–91.

Zhao X, Kong X, Hua Y et al. Medium optimization for lipid production through co-fermentation of glucose and xylose by the oleaginous yeast *Lipomyces starkeyi*. Eur J Lipid Sci Technol 2008;110:405–12.

Zhou YJ, Buijs NA, Zhu Z et al. Production of fatty acid-derived oleochemicals and biofuels by synthetic yeast cell factories. Nat Commun 2016;7:11709.

Zhu Z, Ding Y, Gong Z et al. Dynamics of the lipid droplet proteome of the Oleaginous yeast *Rhodosporidium toruloides*. Eukaryot Cell 2015;14:252–64.

